# Light dynamically regulates growth rate and cellular organisation of the Arabidopsis root meristem

**DOI:** 10.1101/353987

**Authors:** Thomas Blein, Jasmin Duerr, Taras Pasternak, Thomas Haser, Thorsten Falk, Kun Liu, Franck A. Ditengou, Olaf Ronneberger, Klaus Palme

**Affiliations:** Institute for Biology II, Molecular Plant Physiology, Faculty of Biology, Albert-Ludwigs-University Freiburg, Schänzlestr. 1, D-79104 Freiburg, Germany; Institute for Computer Science, Image Analysis Lab, Albert-Ludwigs-University Freiburg, Georges-Köhler-Allee Geb. 52, D-79110 Freiburg, Germany; BIOSS Centre for Biological Signalling Studies, Albert-Ludwigs-University Freiburg, Signalhaus, Schänzlestr. 18, D-79104 Freiburg, Germany; Freiburg Centre for Advanced Studies (FRIAS), Albert-Ludwigs-University Freiburg, Albertstr. 19, D-79104 Freiburg, Germany; Center for Biological Systems Analysis (ZBSA), Albert-Ludwigs-University Freiburg, Habsburger Str. 49, D-79104 Freiburg, Germany

**Author notes:** Current address: Institute of Plant Sciences Paris Saclay IPS2, CNRS, INRA, Université Paris-Sud, Université Evry, Université Paris-Saclay, Orsay, France.

## Abstract

Large-scale methods and robust algorithms are needed for a quantitative analysis of cells status/geometry *in situ*. It allows the understanding the cellular mechanisms that direct organ growth in response to internal and environmental cues. Using advanced whole-stack imaging in combination with pattern analysis, we have developed a new approach to investigate root zonation under different dark/light conditions. This method is based on the determination of 3 different parameters: cell length, cell volume and cell proliferation on the cell-layer level. This method allowed to build a precise quantitative 3D cell atlas of the Arabidopsis root tip. Using this approach we showed that the meristematic (proliferation) zone length differs between cell layers. Considering only the rapid increase of cortex cell length to determine the meristematic zone overestimates of the proliferation zone for epidermis/cortex and underestimates it for pericycle. The use of cell volume instead of cell length to define the meristematic zone correlates better with cell proliferation zone.

## Introduction

Understanding the cellular mechanisms that direct organ growth in response to internal and environmental cues is a major challenge for plant biology (Sozzani and Iyer-Pascuzzi, 2014; Satbhai et al., 2015). Dynamic patterning modules are a key for understanding development. Although numerous studies have provided insight into the regulatory networks that direct organ growth and shape, surprisingly little is known about the precise geometry of the Arabidopsis root and how organs develop in three-dimensional space. Rolland-Lagan et al. (2003) and Coen et al. (2004) determined parameters to characterise plant development in space and time. Hence for each growing region and point in time — the growth rate (the rate of increase in size), the degree of directional growth and the overall form relative to an underlying coordinate. As these parameters continuously changes in space and time, it is essential to determine the quantitative contribution of each of these parameters experimentally. Such data — given in terms of geometric information about cell positions, dimensions and shape — are crucial if we are to build growth models that accurately describe each cell within an organ with respect to their position and in response to signals and signalling pathways (Bao et al., 2006; Long et al., 2009; Thomas and White, 1998). The lack of quantitative methods appropriate for plant organ analysis meant that detailed studies on growth patterns at cellular resolution remained elusive (Baldazzi et al., 2012). Image-based methods have been developed for *in planta* cytometry (Fernandez et al., 2010; Federici et al., 2012; Kierzkowski et al., 2012; Montenegro-Johnson et al., 2015; Bassel et al., 2014). However, despite significant progress in robust image segmentation and tissue reconstruction, no attempt has yet been made to build a plant tissue atlas at cellular resolution and relate this atlas mechanistically to geometric changes caused by external signals of the surrounding environment and the ensuing changes in development. In our investigation we used Arabidopsis roots, as a simplest root model with only 1 cortex layer.

One important question in root plant biology is root zonation. The growing part of the root can be divided into different longitudinal zones or domains. As there is no standard terminology for these zones, we will adopt here the proposal of Ivanov and Dubrovsky (Ivanov and Dubrovsky, 2013; Pacheco-Escobedo et al., 2016). They distinguished a proliferation domain (PD), a transition domain (TD) and a cell elongation zone (EZ). The PD and TD form the root apical meristem (RAM). Other authors partition the root tip in a meristematic zone, elongation zone and differentiation zone (Huang and Schiefelbein, 2015). However, a clear definition and strict rules/algorithms for their measurement are missing. For example, Romero-Arias et al. (2017) determined root zonation based only on visual cortex cell length, without considering other cell files and without robust algorithms. Yang et al. (Yang et al., 2017) determined the meristem length using only the elongation of one unique cortex linearity per root.

Arabidopsis roots consist of several nested, concentric cell layers: the outer layer of epidermal cells surrounds the ground tissues of cortex and endodermis, which in turn enclose the stele with pericycle and vascular phloem and xylem and procambial cells (Dolan et al., 1993). All cell files of the primary root originate from slow-dividing stem cells at the tip of the root apical meristem followed by further rapid cell divisions in the meristematic zone. Accordingly, cell proliferation in the meristematic zone is crucial for root development. In the root tip, initial cells are under the control of the quiescent centre (QC) (Dolan et al., 1993; Weigel and Jürgens, 2002; Sarkar et al., 2007). They form the apical meristem where new cells are produced that differentiate into a range of cell types with different fates, and subsequently pass through transition zones and expand into mature cells (Dolan et al., 1993; Dolan and Roberts, 1995; Scheres et al., 1995, 1996). Each concentric cell layer performs unique biochemical reactions guiding development; hence the generation of a cell atlas will contribute to a better understanding of distinct cellular machineries controlling differential growth between cell layers. Growth consists of cell proliferation and cell elongation, which occur in different zones along the root tip. Cell divisions occur in the meristem zone proximal to the quiescent centre. Cells from this zone then feed into the elongation zone, where cell division ceases and elongation begins. The differentiation zone follows, where cells fully elongate and mature according to their final fate determination. In this region, specialized epidermal (trichoblast) cells develop root hairs. Short, morphologically distinct transition areas separate these zones (Verbelen et al., 2006; Ivanov and Dubrovsky, 2013). Recently several algorithms were proposed based on direct measure of proliferation events in all cell files in Arabidopsis (Lavrekha et al., 2017) and proliferation in combination with DNA replication in tobacco (Pasternak et al., 2017).

Root growth is regulated by both light and temperature, which are arguably the most important environmental signals affecting plant growth and development (Forde, 2009). This plastic growth response was recently shown to depend on phloem transport of photosynthesis-derived sugar, an inter-organ signal from the cotyledons, into the energy consuming sink at the root tip in order to drive cell production, metabolism and root elongation (Kircher and Schopfer, 2012). In order to achieve a quantitative understanding of how root cells acquire their fates in the presence or absence of light, we applied the intrinsic root coordinate system (iRoCS) image analysis pipeline that enables direct quantitative comparison between root tips at single cell resolution (Schmidt et al., 2014). iRoCS allows to monitor subtle changes in cell division patterns within the root apical meristem (RAM) and precisely measure cell geometry of each root cell. Using this tool, we present a cell atlas of the *Arabidopsis* root tip where we first allocate all cells to specific layers and root zones and, second, quantitatively measure their geometric cell dimensions in 3D. So far root zones were defined based on the cell length only (but not cell proliferation) as a marker of cell status. However, cell-length represents only one cellular dimension and therefore may not adequately reflect cell growth. We show that defining meristematic zone using cortex cell-length significantly overestimates the actual zone length when comparing to cell division events distributions. However, when we define the meristem length for each cell layer using their respective cell volume this correlation is significantly improved. To support this observation, we compare root zone quantification based on cell length and cell volume for all cell layers. The application of different light-dark regimes shows that each cell layer has a characteristic cell shape in the transversal section, which remains unmodified under different conditions tested. Alongside these new findings, we provide a robust algorithm for root zone-definition based on cell volume and successfully relate dynamic changes in the cellular geometry under different dark-light regimes to root growth.

### SIGNIFICANCE STATEMENT

A 3D digital atlas of *Arabidopsis thaliana* roots was built from large scale image analysis using the iRoCS Toolbox. The atlas allows to quantitatively analyse the Arabidopsis root apical meristem based on gradients of cell volume, cell elongation and cell proliferation along the root axis. Our analysis shows that changes in cortex cell length do not adequately reflect the length of the proliferation zone. Instead we propose a robust algorithm to define root zonation based on cell volume that correlates well with cell proliferation.

## Results

### Root growth rate correlated with dark-light regime

It was shown that root growth is rapidly and reversibly repressed in the absence of light (Kircher and Schopfer, 2012). Therefore, we applied different light/dark regime to investigate in detail the structure of the RAM under different growth conditions (Figure 1A, B). When comparing seedlings grown in the light or dark we observed no differences in growth rate for the first day after germination since at this stage the majority of the energy required for root growth is provided by the seed reserves. In the presence of light, the growth rate progressively increased up to 6 mm day^−1^ after five days (5dL). However, two days after transfer to the growth room in darkness, root growth rate decreased dramatically until it stopped completely five days after transfer. When light was switched off only after three days (3dL1dD), root growth rate decreased rapidly and after two days of darkness (3dL2dD), it became comparable to that of dark grown seedlings (5dD). When light was turned back on (3dL1dD1dL), growth was restored to a rate comparable to that of a root growing in constant light.

**Figure 1:**
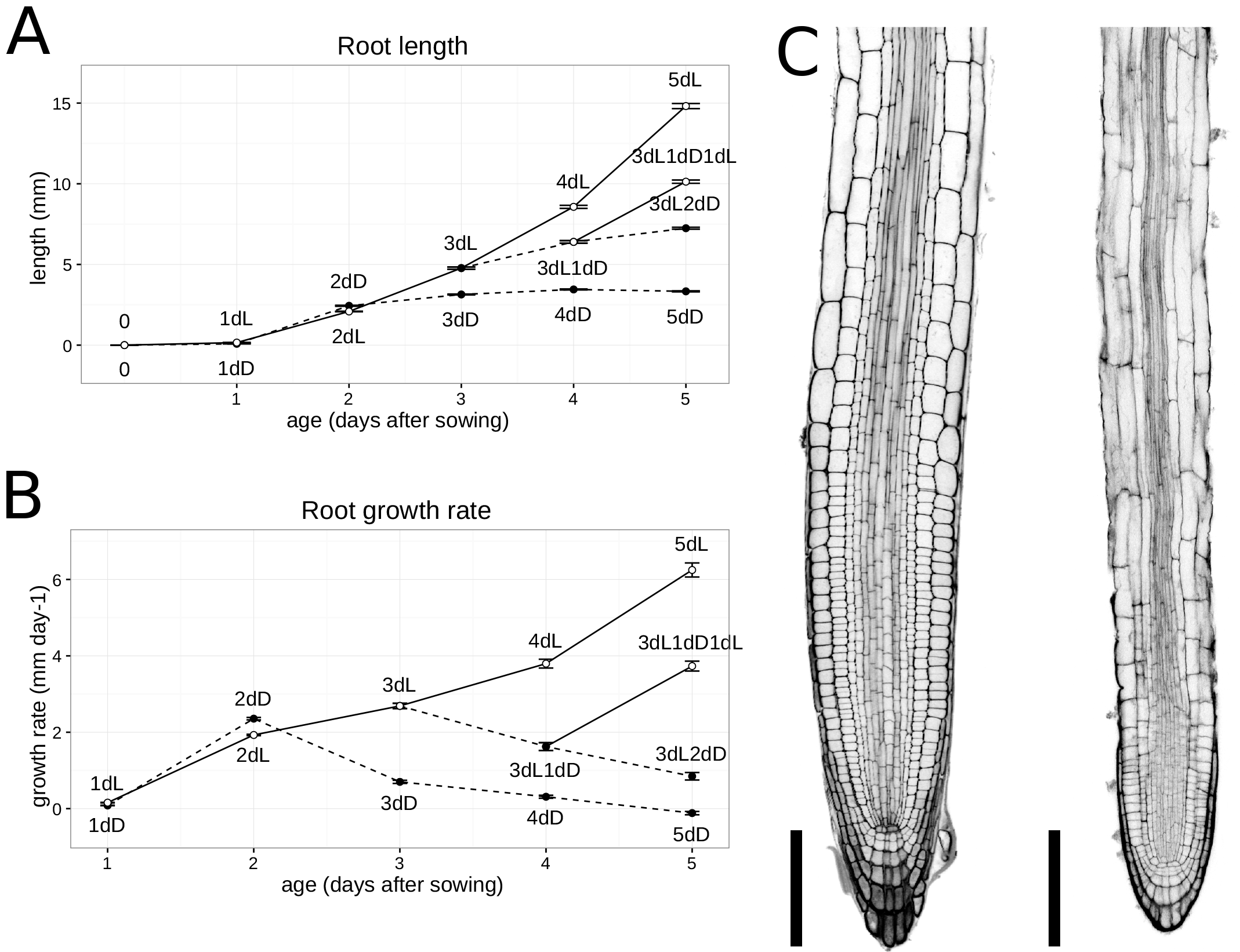
The influence of light on root growth. (A), (B) Influence of light on global root growth. (A) Main root length and (B) main root growth rate according to age and light condition. Values are mean values and error bars indicate SEM. Sample size n = 30 roots for each variant. (C) Morphological appearance of longitudinal sections of root tips grown for 5 days in light or in darkness. Bar = 100 μm. Ages and light conditions are depicted by three characters for each variation: “xdY”, “x” is the number of days the plants stay in that condition, “d” means days and “Y” is either “L” for continuous light or “D” for continuous darkness. In case of modifications of light conditions during the life of the plant this scheme is repeated. For example a plant that first stays three days in light and then two in darkness is labelled as “3dL2dD”.

As expected, the cellular structure of the root tip was clearly different between the two extreme growth conditions (5dL *versus* 5dD, Figure 1C). However, despite changes on the root tip architecture, the overall root structure remained the same: all cell layers are present.

Therefore, varying the light regime can serve as a synchronised plant developmental trigger to understand how root growth is impacted at cellular resolution in response to environmental cues.

### The cellular organisation of the root tip

The three-dimensional shape of an organ is determined by the number, size and organisation of its cells. Growth depends on the cell proliferation and cell elongation. In order to estimate quantitatively how change in root growth is impacted at the cellular level, we extracted features of the proliferating and elongating cells such as the presence of a metaphase plate as marker of cell division and changes in cell shape to display alterations in cell dimension (Figure 2). In the root tip, cells have a typical parallelepiped shape oriented along the root axis for inner cell layers and perpendicular to the root axis for trichoblast cells. Therefore, in addition to their volume, we determined three additional parameters for each cell: their length (along the root axis), thickness (in the radial direction of the root) and width (perpendicular to the root diameter) (Figure 2C).

**Figure 2:**
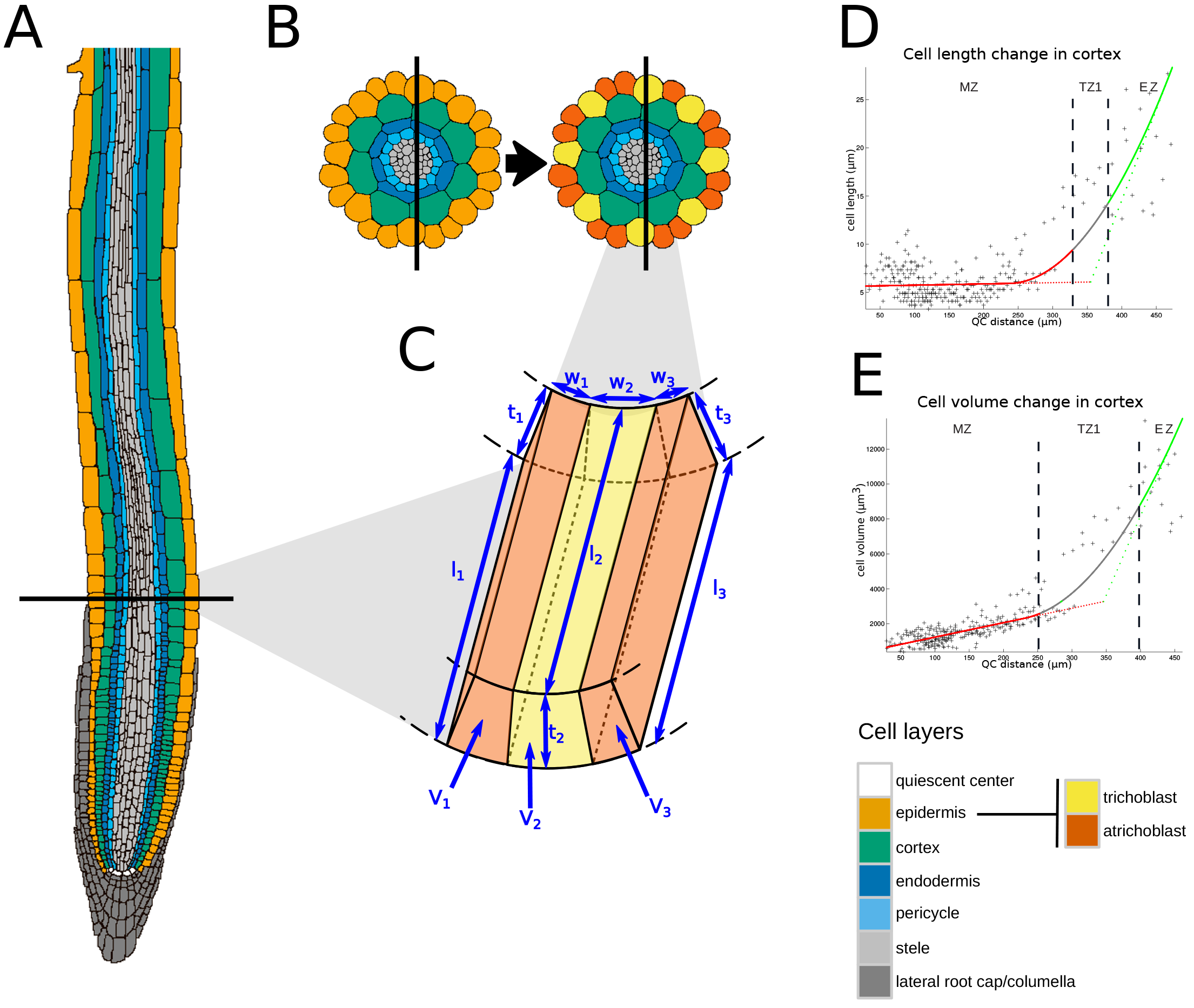
The cellular organisation of the root tip. (A) The layer organization of an A. thaliana root tip. Different colours depict different cell layers (see colour legend at the bottom right of the figure); grey tones indicate cell layers that have not been taken into account in this study. (B - C) Determination of trichoblast and atrichoblast cell types of epidermal cells shown on transversal sections of the primary root (A). Epidermal cells (light orange in B left) are classified according to the number of neighbouring cortex cell files: trichoblast cells (yellow in B right) touch two cortex cell files while atrichoblast cells (dark orange in B right) touch only one cortex cell file. (C) Parameters extracted for each cell of the root. As depicted for three epidermal cells (see (A) and (B)), the cell length (lx), cell width (wx), cell thickness (tx) and cell volume (Vx) were extracted. It should be noted that the orientation of cell width and thickness vary; the thickness passes off orthogonally, the width tangentially towards the root’s circular cross section. (D) Extraction of root zonation according to cell length in the cortex of one root. A continuous function f(x) is fitted (continuous line) to the observed cell position and cell length (each cell as “+”). This model is then simplified to a piece-wise linear function fl(x) (dotted line). From this model the developmental zones are extracted. Extracted zones are indicated by colours (meristematic zone MZ, elongation zone EZ). Grey indicates transitions between zones (TZ1). (E) Extraction of root zonation according to cell volume in the cortex of one root. The similar approach as for cell length is used. The zonation is established in the same root as in (D).

We then used the iRoCS pipeline (Schmidt et al., 2014) to locate each cell within its layer and determine its distance from QC (Figure 2). It is known that epidermal trichoblast and atrichoblast cells distinguish according to the number of adjacent cortex cell files (Galway et al., 1994; Berger et al., 1998). By integrating this feature into the iRoCS pipeline, we were able to automatically allocate all epidermal cells as trichoblast or atrichoblast cells (Figure 2B, C). This allowed the automatic allocation of each cell in its specific root cell layer.

### Determining root zonation

The root is constituted of cells with individual fates and functions, which are arranged in different development zones along the root axis. Classically, the three development zones of the root (meristematic, elongation and differentiation zones) are determined using the cell length as a proxy of cell size (Ivanov and Dubrovsky, 2013). For example, the boundary between the meristematic and elongation zone is usually determined as the point where the length of cortex cells begins to increase rapidly (Casamitjana-Martínez et al., 2003).

First, we reconstructed the zonation of several roots of light grown plants (n=16) from 3D image stacks using a model fitted on cell length to precisely define the different root zones with respect to the cellular geometry (Figure 2D; see Supplemental Figure 1A). We used a mathematical model that determines the borders of the different zones as follows: (1) in the meristematic zone (MZ) cell length is on average constant; (2) in the elongation zone (EZ) cell length increases linearly; (3) in the differentiation zone (DZ) cell length reaches a plateau. As illustrated in Figure 1D and 1E, the transition between these main zones is made up of two distinct zones. The first transition zone (TZ1) is located between the MZ and the EZ (Figure 2D), whereas the second transition zone (TZ2) is between the EZ and the DZ (Supplemental Figure 1A).

The precision of the average cell length determination decreases as the cell size increases due to decrease in sampling: as cell length increases less cells are present at a given point to estimate the average size at that point. We were able to determine the zone borders in small cell zones at a high precision for the MZ, TZ1 and the beginning of the EZ. Due to rapid cell size increase, our analysis was less precise for borders at the end of the EZ and for the TZ2 and DZ (see Supplemental Figure 1A). Therefore, the analysis was focused on the MZ and TZ1.

Extracting the full 3D cellular structure of the root gave us access in addition to cell length, to cell volume that can be used as another estimation of cell size. We adapted our model to predict root zonation using cell volume (Figure 2E; see Supplemental Figure 1B). A strong correlation was detected between the predictions of the MZ size according to cell volume and according to cell length for the fast growing roots, grown under 5 days of continuous light (Figure 3A). This linear relationship is still true for other light conditions (see Supplemental Figure 2A, C, E, G, I). In most of the case the MZ length prediction is bigger when calculated according to cell length than to cell volume for fast growing roots (Figure 3B). This tendency is conserved for the different light conditions (see Supplemental Figure 2B, D, F, H, J). However, the difference of the prediction is decreasing with the decrease of growth rate with no difference for non-growing roots (see Supplemental Figure 2).

**Figure 3:**
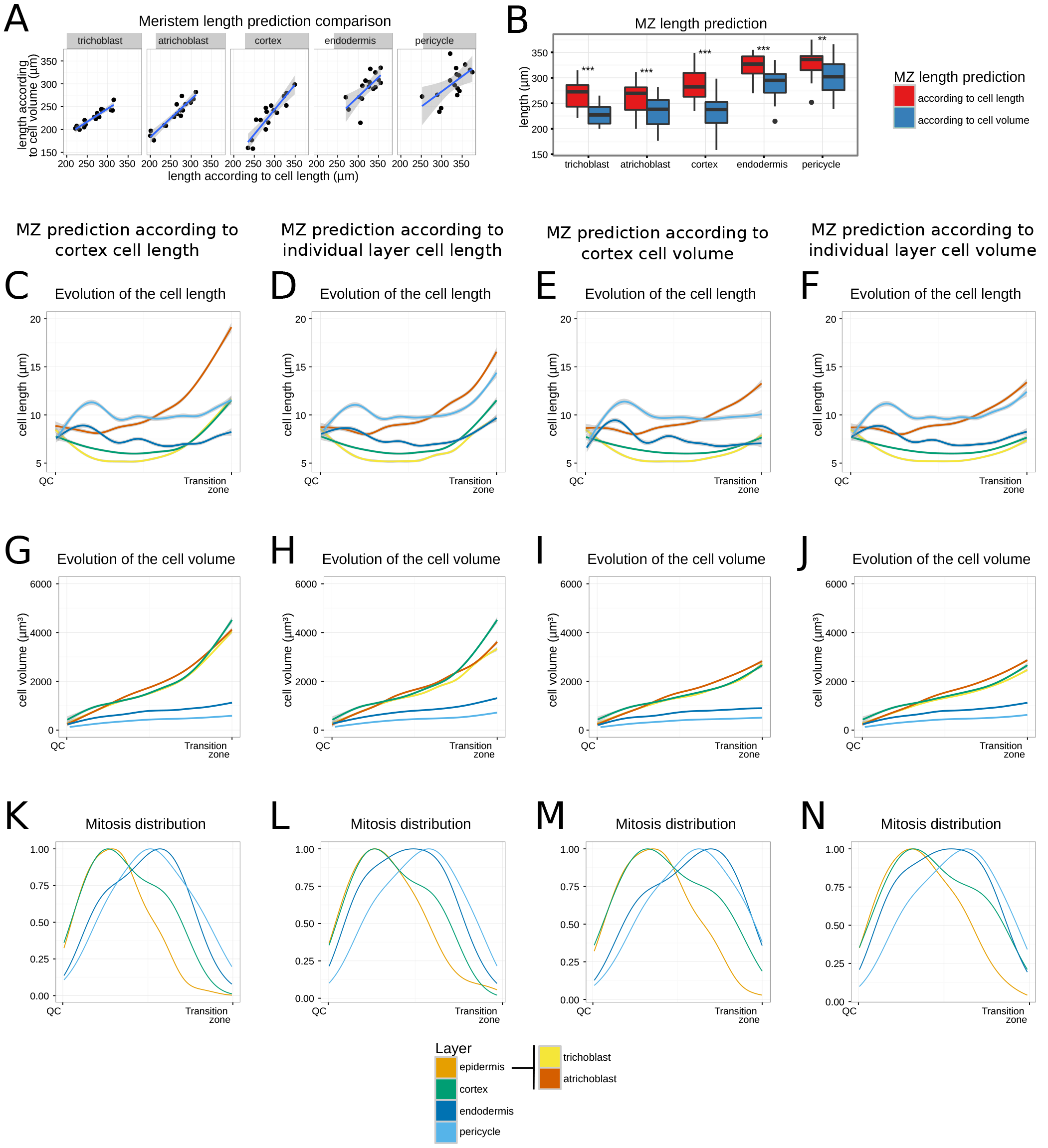
The influence of the parameter used for meristem length prediction for fast growing roots. (A) Relation between the prediction of meristem length according to cell length and according to cell volume for each cell layer and root grown for 5 days in light. Curves represent a linear regression fitted to the data. The grey shadow indicates SEM. (B) Distribution of meristem length prediction according to cell length and cell volume for root grown for 5 days in light. Stars indicate significant differences between the two predictions from a Wilcoxon Rank Sum test: ** P<0.01, *** P<0.001. (C - F) Evolution of cell length along the root axis according to different meristem prediction methods. Curves represent a smooth representation of the parameter fitted to the data using generalised additive models. The grey shadow indicates SEM. (G - J) Evolution of cell volume along the root axis according to different meristem prediction methods. Curves represent a smooth representation of the parameter fitted to the data using generalised additive models. The grey shadow indicates SEM. (K - N) Distribution of mitoses along the root axis according to different meristem prediction methods. (C, G and K) Prediction of the meristem according to cell length in cortex cell layer. (D, H and L) Prediction of the meristem according to cell length in individual layer. (E, I and M) Prediction of the meristem according to cell volume in cortex cell layer. (F, J and N) Prediction of the meristem according to cell volume in individual layer. Data extracted from roots marked cell walls (n = 16 roots) and from roots marked cell nuclei (n = 19 roots).

Despite the highly altered cellular morphology of the non-growing root, after five days of darkness, our zonation model is still able to predict the different zones (see Supplemental Figure 1C and D).

### Influence of MZ definition on cellular characteristic distribution of the fast growing root tip

We wondered how the different zone prediction models may influence the distribution of cell sizes and mitosis in the MZ depending on: i) the prediction according to cell length or cell volume ii) the prediction using only cortex or each layer individually. We used the border of MZ to normalize the cell position along the root axis, thereby allowing the precise longitudinal placement of each cell and the comparison of different roots, even in cases where the zones displayed large variations in length.

When using only the cortex as a determinant of MZ, cellular parameters clearly showed that it does not fit for all the layers (Figure 3C, G, K). For example, we clearly see a strong increase of cell length in trichoblast and atrichoblast cell layers that would fit better in TZ1 than MZ. In contrast, the inner cell layers do not show the small increase of cell length at the end of the predicted MZ (Figure 3C). It is even clearer for mitosis distribution, where a clear drop of mitosis in epidermis before the end of the MZ could be seen (Figure 3K).

When predicting the size of the meristem for each layer individually, the shape of the distributions for cell length is more homogeneous between the layers (Figure 3D) and the drop in mitosis distribution for epidermis is not as strong (Figure 3L). However, using cell length to predict MZ border, showed a strong difference of cell volume evolution between inner cell layers and outer cell layers (Figure 3G, H). The cell volume of inner cell layers grows linearly in MZ while for the outer cell layers, a strong increase in progression appeared at about 75% of MZ length.

Using volume for MZ prediction homogenised even more the distribution pattern between the different cell layers. All cell layers showed a slight increase of cell length at the end of the MZ (Figure 3F). No volume progression rupture was seen (Figure 3J) and the MZ stopped roughly when the number of mitosis decreased, with a distribution peak closer to the middle of the MZ, except for epidermis which seems to be cut too short (Figure 3N). We concluded that cell volume seems a better parameter to predict MZ limits than the classically used cell length. If determining the MZ with only the cortex cell volume the shape of the distribution for cell length and cell volume is comparable as with individual cell layers (Figure 3E, I). However, the mitosis distribution for endodermis is clearly shift in direction of the shootwards border of the MZ (Figure 3M). Therefore, from now on the MZ definition used in this study will be based on the prediction according to cell volume and defined individually for each cell layer.

### The cellular structure of a fast growing root MZ

It is an accepted view that in order to preserve structural organ integrity, the global growth rate of the different layers should be identical (Band et al., 2012). However, it is also accepted that—during organ development—cell proliferation and expansion may differentially contribute to growth and change in growth rates of different cell layers. In order to test whether the different root cell layers grow homogeneously, we investigated whether changes in the number of cells and/or cell size contribute to the observed differences in zone sizes. Since the different cell layers do not have the same number of cell files (Dolan et al., 1993) we calculated the number of cells per cell file. Hence, by comparing them to the average cell length we would see a direct anti-correlation between the cell number and the average cell length.

Two groups of layers can be defined: the first group (trichoblast, cortex and endodermis cell files) consisted of cell files with about 40 cells with the smaller average cell length (5-7μm long) in MZ. The second group (atrichoblast and pericycle cell files) consisted of about 30 longer cells (10μm long) (Figure 4A, B).

**Figure 4:**
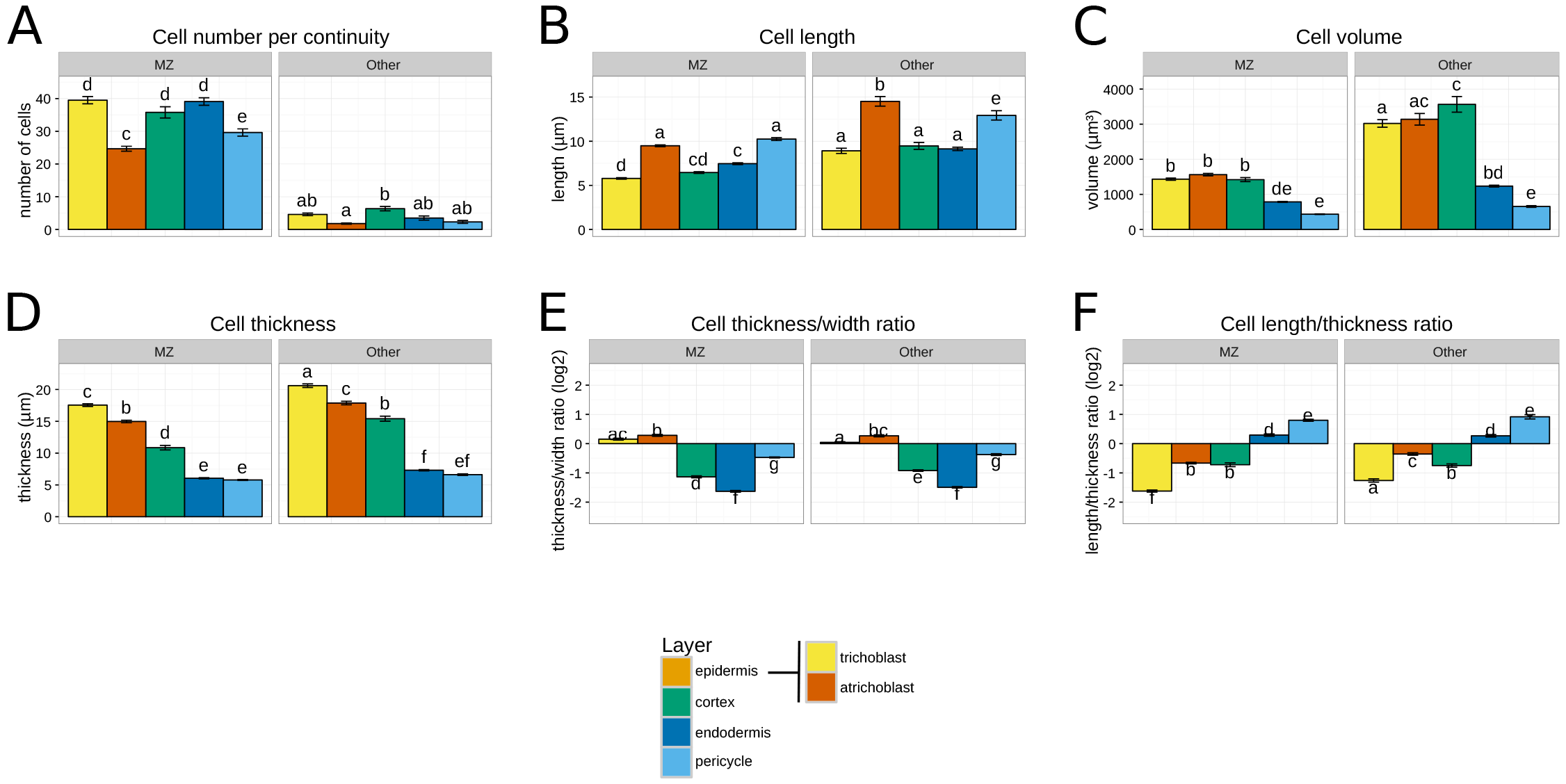
Average cellular geometry of the growing root. (A) Number of cells per cell file according to cell layer and developmental zone. (B - D) Average cell dimensions according to the layer and developmental zone: (B) length, (C) volume, (D) thickness. (E - F) Average cell shape according to layer and developmental zone: (E) average ratio between cell thickness and cell width, (F) average ratio between cell length and cell thickness. The developmental zone is marked as MZ when both predictions (according to cell length and according to cell volume) predict the cell to belong to meristem. It is marked as other when only the prediction according to cell length predicted it as belonging to meristem. Values are the means and error bars SEM. Different letters display significant differences between the layers by a Tukey’s test at a 95 percent confidence level. Data extracted from roots marked cell walls (n = 16 roots).

As seen before, cell length is not homogeneous in MZ (Figure 3F). We can see a slight diminution of cell length for each cell layer that correlate nicely with the peak of mitosis in the MZ (Figure 3F, N). More precisely we can highlight three different patterns in the evolution of cell length along root axis (Figure 3F). First, atrichoblast cells have a short decrease of division at the rootward border of the MZ and then constantly increase their cell length until leaving the MZ. Second, for trichoblast and cortex cell layers, we can observe a continuous decrease of cell length until the middle of the MZ and then a smooth increase up to the shootward border of the MZ. Third, endodermis and pericycle cell layer show a small increase in cell length at the rootward border of the MZ and then the decrease linked with mitosis maximum. Indeed for these cell layers the maximum of division is positioned more shootward. Interestingly, independent of the cell layer, cell length at the rootward border of the MZ are the same, while at the shootward border the layer can be classified in the same two categories as before: atrichoblast and pericycle cell layers having longer cells (length around 12μm) than trichoblast, cortex and endodermis cell layers (length around 8μm). These data lead us to conclude that cell length follows a common trend along the root axis for all cell layers, where the average cell length varies mainly according to the distribution of mitosis.

The determination of the cell volume provided another classification: the outer cell layers (trichoblast, atrichoblast and cortex) have bigger cells than inner cell layers (endodermis and pericycle) (Figure 4C). When comparing average cell thickness, we see even more differences between outer cell layers with a global tendency to decrease cell thickness while going inside the root (Figure 4D). These measures showed a constant increase in thickness inside the meristem for each layer stronger for the outer cell layers (Figure 3J; see Supplemental Figure 3B).

Next, analysing the isotropy of the cell showed that each cell layer has a specific ratio between its dimensions. For example cortex, endodermis and pericycle cells are wider than thick, while trichoblast and atrichoblast cells are as wide as thick and are therefore isotropic (Figure 4E). This relation is conserved all along the MZ and is characteristic of each cell layer (see Supplemental Figure 3F). In other words, cells are growing as fast in thickness as in width. The isotropy along the root axis, illustrated by the length/thickness ratio, is also characteristic of each cell layer (Figure 4F). However, due to the stability of the cell length in the MZ, this ratio decreases progressively in the MZ while moving shootward. Cell thickness increases faster than cell length with a small inversion at the shootward border of the MZ, where cells start to grow faster along root axis, *i.e*. elongate, than along root section (see Supplemental Figure 3D). As a result, the increase of cell size in MZ as described by the progressive increase of cell volume along the MZ is mainly due to an increase in cell diameter in the MZ and then to cell length at the shootward border of the MZ.

The differences between the MZ prediction according to cell volume and according to cell length allowed us to define an additional zone that we named “other”. This zone is present only in growing roots (5dL, 3dL1dD and 3dL1dD1dL; see Supplemental Figure 2) but disappears when root growth stops (5dD and 3dL2dD; see Supplemental Figure 2). In fast growing roots, this zone is constituted of few cells from 1 to 8 cells in respectively atrichoblast and cortex layers (Figure 4A). There, cells are in average bigger in all directions compared to those from MZ (Figure 4B-D) but conserve the same shape as their respective MZ counterparts (Figure 4E, F). This corresponds to a zone located between the end of proliferation zone and the area of fast cell growth.

### The cellular structure of the arrested root

Next, we investigated the influence of root growth on cell geometry in tips of roots that stopped growing after 5dD. Our image analysis pipeline allowed the direct comparison of roots with different morphologies (Figure 2D and E; see Supplemental Figure 1C and D), revealing drastic changes in length of the MZ in every layer after 5dD (Figure 5A). This enabled us to investigate whether the reduction of MZ lengths was caused by a reduction in cell proliferation or cell elongation.

**Figure 5:**
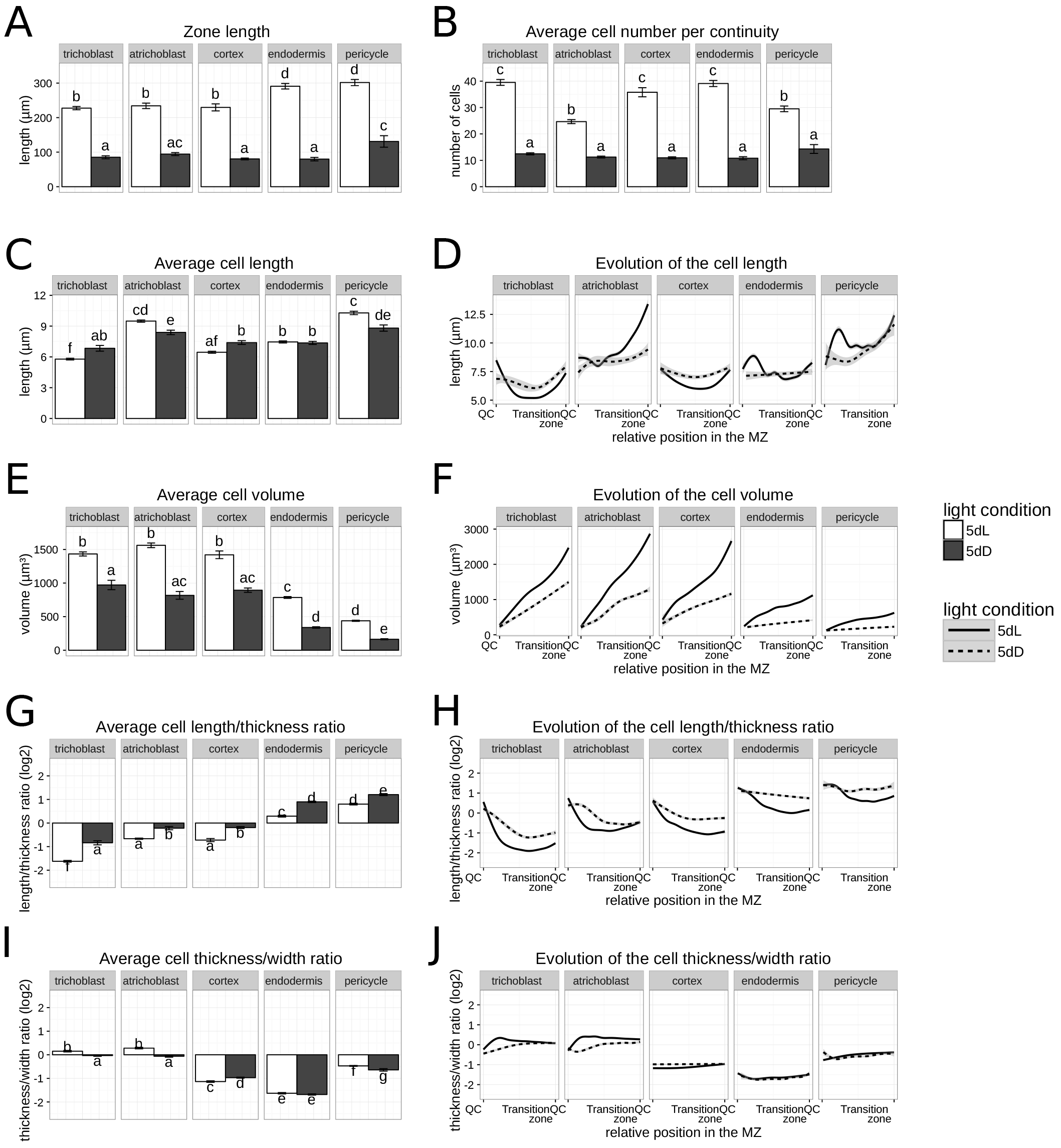
The influence of long darkness treatment on the cellular geometry of the MZ. (A) Meristematic zone length according to cell layer and light regime. (B) Cell layer- and light regime-dependent classification of the number of cells per cell file in the MZ. (C - J) Average and evolution of cell parameters along the meristem according to cell layer and light regime: (C - D) cell length, (E - F) cell volume, (G - H) ratio between length and thickness, (I - J) ratio between thickness and width. (C, E, G and I) Average cell dimensions in the MZ according to cell layer and light regime. Values are mean values and error bars, the SEM. Stars indicate significant differences between the two growing conditions from a Student t-test: * P<0.05, ** P<0.01, *** P<0.001. (D, F, H and J) Evolution of cell length along the root axis according to cell layer and light regime. Curves represent a smooth representation of the parameter fitted to the data using generalised additive models. The grey shadow indicates SEM. Data extracted from roots with marked cell walls (n = 16 roots for 5dL and n = 15 roots for 5dD).

In roots that stopped growing no cell division was detected (see Supplemental Figure 4). In these roots, the reduction of MZ length strongly correlated with a decrease in cell number which dropped to about 10 cells per cell file (Figure 5B). A small difference in average cell length was observed between the non-growing root tip and the fast growing one (Figure 5C). The analysis of cell length distribution along the MZ showed a different profile compared to the fast growing root: cell length was almost constant throughout the MZ with a slight increase at the end of this zone (Figure 5D). The decrease of cell length that was linked with the position of high mitosis density in fast growing roots is strongly reduced like in cortex cell layer or disappears completely for the other cell layer (Figure 5D). This reinforces the correlation between the presence of cell division and the specific profile of cell length distribution observed in fast growing roots.

We then investigated the other cell parameter measurements in the dark grown root. The cell volume was strongly decreased for all cell layers (Figure 5E). As in fast growing roots cell volume in arrested roots increased in the MZ with the distance from the QC, though with a lower rate than light-grown plants (Figure 5F). The cell length/thickness ratio was increased in non-growing roots highlighting the higher contribution of the cell length in the increase of cell size in the non-growing roots compared to fast growing roots (Figure 5G,H). The ratio cell thickness /cell width is almost constant between the two growth regimes (Figure 5I). The differential growth observed is therefore not modified except in the rootward border of the MZ for trichoblast and atrichoblast cell layers (Figure 5J). In arrested roots, cells are growing less but more homogeneously in all dimensions when compared to cells in fast growing roots.

### Growth reduction correlates with a decrease in cell size and number in the MZ

To understand the cellular basis of growth reduction, we investigated the proliferation and the elongation of cells during the transition from a high to low growth rate. For this, seedlings grown in light for three days were transferred to darkness for one or two days (3dL1dD and 3dL2dD). Compared to roots growing under continuous light, the growth rate was reduced by 50 percent after 24 hours in darkness. An additional day in darkness decreased the growth rate by a further 50 percent (Figure 1A, B).

Roots growing at a low rate have a shorter MZ than fast growing roots (Figure 6A). No cell divisions were observed in both conditions (see Supplemental Figure 2A, B) and the reduction of MZ length correlated with a decrease in the number of cells (Figure 6B). Despite the differences in growth rate after one or two days in darkness, no other differences in MZ length or number of cells were observed between these two treatments (Figure 6A, B).

**Figure 6:**
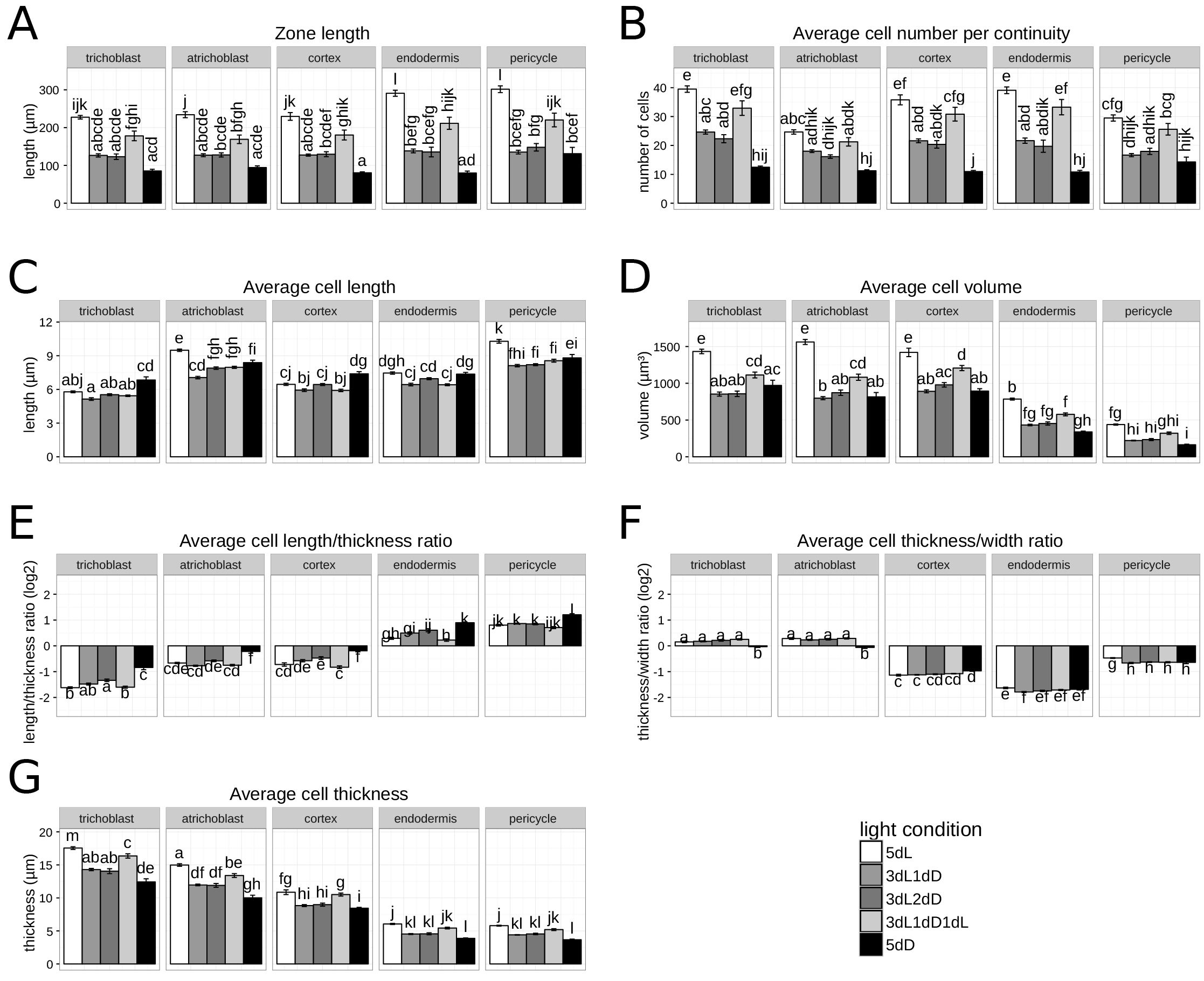
The influence of growth rate on the average cellular geometry in MZ. (A) Meristematic zone length according light regime. (B) Number of cells per cell file according to light regime. (C - G) Average cell parameters in MZ according light regime: (C) cell length, (D) cell volume, (E) ratio between length and thickness (F) ratio between thickness and width (G) thickness. Values are the means and error bars SEM. Different letters display significant differences between light regimes from a Tukey’s test at 95 percent confidence level. Data extracted from roots with marked cell walls (n = 16 roots for 5dL, n = 16 roots for 3dL1dD, n = 17roots for 3dL2dD, n = 15 roots for 3dL1dD1dL and n = 15 roots for 5dD).

Compared to fast growing roots, cell length, thickness and volume were decreased in roots growing slowly (Figure 6C, D, G). No changes in cell shape were visible except for the cell length/width ratio that was slightly increased for all cell layers compared to fast growing roots (Figure 6E-F). In slow growing roots, cells had a similar distribution along the entire MZ when compared to cells of more rapidly growing roots (Supplemental Figure 5A-E). The decrease in cell length in the middle of the MZ was less pronounced after one day of darkness than in fast growing roots. An additional day in darkness decreased this difference further (Supplemental Figure 5A). These growth conditions represent intermediate stages between the fast growing (5dL) and the arrested roots (5dD) in the shape of cell length distribution along the root axis. It suggests that root growth does not stop abruptly but occurs gradually through changes of cellular features. This has to be related with the absence of cell division in the middle of the MZ for these two conditions (see Supplemental Figure 4).

### Growth recovery of slow growing roots restores both cell proliferation and cell size

We then analysed cellular changes occurring during root growth recovery induced by one day of light after one day of darkness (3dL1dD1dL). Under this condition, we observed almost a doubling of the growth rate compared with the previous time point (3dL1dD) (Figure 1A, B). When root growth resumed, the MZ length increase correlated with a restoration of cell division in the MZ (Figure 6A; see Supplemental Figure 4). The mitotic rate of 1.9 percent was similar to fast growing roots. However, the number of cell increased but not to the same extent to that of fast growing roots (5dL) (Figure 6B).

A determination of the distribution of cell length along the MZ showed the same pattern for all layers as it was observed in fast growing roots (see Supplemental Figure 5A). On average, cell length did not differ significantly from slower growing roots (Figure 6C) but cells were longer at the end of the MZ (see Supplemental Figure 5A). This further emphasized the differences between the middle and the end of the MZ (see Supplemental Figure 5A). However, at the end of the MZ, cell length did not reach the length of the fast growing roots of the same age (see Supplemental Figure 5A). Further, the thickness and volume of cells increased in comparison to that roots growing at a slower speed (Figure 6D, G; see Supplemental Figure 5B, C). As was previously observed, no major change in cell shape was noted (Figure 6E-F; see Supplemental Figure 5D-E). Altogether, these data suggest that when the root growth resumes the mitotic activity restarts first, which then determines the number of cells in the different zones. Subsequently, cells globally expand in all dimensions.

## Discussion

### Determination of the longitudinal root zonation pattern

Root growth and development are dependent on combined activity of two processes that are closely linked: cell division and cell elongation. Precise quantitative characterization of both processes in roots is thus essential for understanding its role during development. The root of *Arabidopsis thaliana* serves as an important model for molecular genetics and cellular studies of plant development. More than 20 years ago simple protocol for investigation of the root zonation has been proposed (Beemster and Baskin, 1998). However, since then several methods have been developed (Truernit et al., 2008; Pasternak et al., 2017) for a more precise analysis of the plant root organization. Despite of these achievements, robust algorithm and basic principles are lacking to define longitudinal zonation of root apices (Ivanov and Dubrovsky, 2013). The classical approach used to estimate RAM length is to use the distance from the QC to the first rapidly elongated cortex cell (Sabatini et al., 1999; Dello Ioio et al., 2007). Recent investigations proposed more complex algorithms for root zonation: based on the cortex cell length distribution along the root axis such as the “multiple structural change algorithm” (Pacheco-Escobedo et al., 2016) or the “Stripflow algorithm” (Yang et al., 2017). Root cap length was also proposed as another criterion to estimate the RAM length (Fendrych et al., 2014). However, none of current investigations distinguished between cell layers and showed meristem length as a unique measurement common to all cell layers. Therefore, only one cell layer is commonly used to determine meristem length, mainly using the cortex cells as a reference.

Here, using quantitative 3D root reconstruction, we analysed the changes in cell volume, cell length and mitosis distribution along the root tip axis for different cell layers: epidermis (trichoblast, atrichoblast), cortex, endodermis and pericycle.

In the MZ, due to cell proliferation, cell size mainly oscillated between two sizes: size after cell division (early G1) and size before cell division (late G2) (Brumfield, 1942; van der Weele, 2003). Previous approaches were using changes in cell length as cell size estimator and stay in 2D. However, root cells are not cylindrical, especially in the outer cell layers. This fact raised the question if cell length adequately reflects cell size. Using 3D imaging gave us the opportunity to test if the volume of a cell can be used as a better estimator of zone size when compared to cell length. Hence, while the classical prediction of MZ length according to cortex cell length (Sabatini et al., 1999) gives overestimation of the MZ length that particularly accentuated in the cortex and epidermis, our data show that using MZ prediction according to volume of the individual cell layers clearly fit better with mitosis distribution.

Cells in the TZ1 progressively increase their elongation rate until they reach the linear elongation rate observed in the EZ. Given these regular elongation patterns, the borders of each zone are computationally accessible despite the variation in cell geometry caused by different growth rates. The definition of the root zones could then be used to normalize the positions of cells within root apices of different sizes and easily compare them.

Although cells of the TZ1 integrate different signals that determine their fate, the morphological specification of this zone is not precisely defined (Baluška et al., 2010). Ivanov and Dubrovsky (2013) defined the TZ1 as a zone with a growth rate similar to that of the MZ but with a low probability of cell divisions. As defined in our study here, this zone corresponds to the portion of the root that would be predicted as in the MZ using cell length but not if using cell volume. This sub-zone is well defined in growing roots or roots where growth is declining, but cannot be determined in a non-growing root. Our analysis used the evolution of the cell volume along the root axis to determine the different root zones and in a second step, the MZ subzones, as a majority of studies do (Ivanov and Dubrovsky, 2013). MZ is generally considered as *de novo* cell production. However, our data show that only a part of the MZ is involved in that process: the cell proliferation sub-zone. Its positions varies within layers: in outer cell layers (trichoblast, atrichoblast and cortex), this zone is shifted rootward with a peak of mitosis activity at 25 % of MZ length, while in inner cell layers the peak of mitosis is localised in the middle of the MZ. This has a direct consequence on the distribution of cell length and volume along the root axis.

### Each cell layer has a characteristic cell shape

The shape of cells in transverse sections is unique and characteristic for each cell layer. Our analysis of root images allowed both the comparison and high resolution phenotyping of different roots and revealed the morphological properties of cells in different zones of roots. This parameter was so far never analysed quantitatively. Our results revealed, for example, that in the cross section of the MZ both the thickness and width of cells decreased from the outer to the inner cell layer. Since root cell layers are concentric, the reduction of cell width correlates well with the decrease in cell layer perimeter. The only notable exceptions were the trichoblast and atrichoblast cells that are smaller in width than the cells found in the cortex. While the cortex consists of 8 cell files, in the epidermis the number of cell files strongly increased to an average of 19 (Dolan et al., 1993). Compared to the cortex, there is a correlation between the reduction of cell width and the increase in number of cell files in the epidermis.

### Evolution of cell characteristics along the MZ

The resulting growth of the different layers is the same due to the physical connection between them (Band et al., 2012). To reach the same layer elongation rate, two different strategies can be assumed: roots either produce many small cells or just a few long cells. Already in the MZ both strategies can be seen in to adjacent cell files. For instance, atrichoblast cells are longer than trichoblast cells though both cell files have a comparable MZ length. This is related to the fact that trichoblast cells have a lower proliferation rate (Galway et al., 1994; Berger et al., 1998). On the other hand, atrichoblast cells also have a higher elongation rate in the MZ when compared to the other layers. This may explain why no significant decrease in cell length can be observed in the middle region of the MZ as seen in the other layers. These data suggest that atrichoblast cells do not only elongate to prepare for division but also increase their average cell length without necessarily dividing.

Despite layer specific cell geometry, our data show that each layer maintains the same progression in size along the growth axis, with a stronger increase of cell volume in the outer cell layers: cell width and thickness increase strongly in the rootward part of the MZ and more slowly in the shootward part to reach a plateau; cell length increases close to the rootward border of the MZ, then decrease due to cell division and finally increase strongly at the shootward border of the MZ, indicating the beginning of the elongation transition; the cell volume increases slowly in the rootward part of the MZ, driven by increase in cell width and thickness, and then increases strongly in the shootward part of the MZ, driven by cell length increase. This allowed for the identification of three sub-zones in the MZ. In the root apex (rootward) oriented sub-zone of the MZ, cell growth was found to be anisotropic in the cortex and epidermis. In this zone cells there displayed a low mitotic index, as they were still under the influence of the QC (Van Den Berg et al., 1997). While the MZ is usually considered as uniform, our study showed that the majority of mitosis occurred in the next sub-zone found around the middle of the MZ in the inner cell layers, while in the outer cell layers the peak of mitotic activity was shifted rootward (fig, 3, K-N). This directly affects the resulting cell length variations, since each division leads to a decrease in cell length in the region where cell division preferentially occurs. In the shootward sub-zone of the MZ, proliferation ceased and cell length slightly increased until cells reached their final size in the MZ: from 10.7 μm in endodermis (SD 3.2 μm) to 14.1 μm in pericycle (SD 4.5 μm). Afterwards, cells left the MZ and started to elongate. Our results suggest that cells grow at a constant rate in the entire MZ since the size differences observed are tightly linked with their proliferation. However, after an extended period of darkness (5 days) during which the root is not growing, the sub-zonation of the MZ is no more visible, due to a complete absence of cell division.

### The change in growth rate is mainly influenced by cell proliferation

To understand the relationship between cellular organization and root growth rate we used our recently developed iRoCS image analysis pipeline and employed its features to study the cellular organization of roots growing at different rates as a result of different light regimes. Our investigations revealed for the first time the detailed three-dimensional cellular organization of the root tip in a quantitative manner.

The easy manipulation of light allows us to switch on and off root growth quantitatively (Kircher and Schopfer, 2012) and very rapidly, in less than 24 hours. Our data show that cell proliferation is the first parameter that is modified in response to changes in the light regime. While a strong variation in cell length is visible in growing roots, this variation is reduced when the growth rate decreases (stop) until it is no longer visible when the root stops growing. But the comparison of average cell length in the entire MZ shows no significant difference. Taken together, these data show that—in order to understand changes in growth rate— it is necessary to analyse changes in cell length depending on the position in the MZ.

The growth rate is proportional to the MZ size, which in turn strongly correlates with the number of cells that constitute it. Indeed, the number of cells is the major factor controlling the size of the MZ. However, cell size, in all directions, is also influenced by the growth rate with thick growing roots and thin non-growing ones. At the end of the MZ cell length is related to growth rate: the more the roots are exposed to light, the longer the cells. This suggests an influence of the cell elongation rate on the EZ on the growth rate.

In the present investigation, we establish for the first time a full three-dimensional atlas of the Arabidopsis root. It allows the monitoring of development of the different cells and their distinct dimensions. Our quantitative analysis highlights the subdivision of the meristem zone in different subzones related to the mitotic activity. In addition, we link these quantitative cell parameters with the root growth rate and how they change in response to light.

## Conclusions

The results presented here show that the iRoCS Toolbox computational root analysis pipeline allows the creation of detailed root maps and provide a robust algorithm for automatic precise root zonation. So far root zonation in majority of the investigation was performed only approximately, based on only single cell continuity of the cortex. It should be noted that iRoCS analysis is based on features that do not require the use of transgenic lines with marker genes, and can therefore be applied to any plants.

## Experimental Procedures

### Plant Material and growth conditions

Arabidopsis thaliana ecotype Columbia (Col-0) was used in this study. Seeds were surface-sterilized and sown on solid ½ Murashige and Skoog medium (Tanaka et al., 2001) with vitamins (Duchefa, M0222) supplemented with 5 mM MES and 1.1 percent agar (Roth). The pH was set to 5.6. After vernalisation for 16 h at 4 °C plants were exposed to white light (100 μmol m ^−1^ s ^−1^) for 4-5 h before transfer to the final growth conditions. Seedlings were grown on plates placed vertically in an incubator and kept at 24°C in continuous white light with an intensity of 100 μmol s ^−1^ m ^−1^. For growth in darkness, the plates were covered with a black tissue and placed in a black box in the same incubator. Days of growth were counted after the transfer from 4°C to the growth chamber. The plates were scanned with a Canon 950 scanner and root lengths were measured using ImageJ.

### Staining

For staining the nuclei, the seedlings were fixed in 2 % percent formaldehyde for 45 minutes, then washed with water, treated with methanol for 20 minutes, rehydrated with water and mounted on microscopy slides with mounting media containing DAPI (DAPI-pro-Gold). The cell wall staining was conducted as in Truernit et al. (2008) with an overnight amylase treatment at 37 °C as previously described (Wuyts et al., 2011). Samples were mounted on slides in mounting solution (80 g of chloral hydrate in 27 ml H_2_O, 3 ml of glycerol) with a 100 μm thick PVC adhesive tap as spacer. Stained roots were recorded using a LSM 510 laser scanning microscope with a Plan-Neofluar 40x/1.3 oil objective. Serial optical sections were reconstituted into 3D image stacks, with a resolution of 0.4 μm (x,y) and a section spacing of 0.4 μm.

### Extraction of cell parameters

After recording, consecutive images were stitched to a total length of 1200 μm from the QC using the XuvTools software (Emmenlauer et al., 2009). After manually indicating the position of the QC, the cell nuclei or cell walls were identified using iRoCS (Schmidt et al., 2014). The layers were determined using the same software for five roots with completely corrected annotations as a training set for the nuclei and 10 roots for the cell shape. Expert corrections were then processed for each automatically annotated root.

### Statistical analysis

All statistical analysis were done using R version 3.2.5 (http://www.r-project.org/) and visualized in the package ggplot2 (Wickham, 2009). The fitting curves represent a smooth representation of the measured parameter fitted to the data using generalized additive models (gam) as implemented in R.

### Zone modelling

For each cell layer and root, the root zonation was determined using three linear models: in the MZ cell length is constant; in the EZ cell length increases linearly; in the DZ cell length is constant (Figure 1D). For the determination of the TZ1, the local average cell length was compared to the two theoretical ones obtained through the linear models of the MZ and the EZ. We determined the TZ1 as the portion of the root where the differences to the two linear models are comparable. For the TZ2, we used the same approach but with the linear models of the EZ and the DZ. By applying the same mathematical determination, root zonation was determined for each individual cell layer (see Supplemental Method).

### Automatic determination of the trichoblast and atrichoblast cell type

For each epidermal cell, the distance to each neighbouring cortex cell in radian was computed according to the φ angle in the iRoCS coordinate system. Cells were then partitioned according to their distance from the epidermal cell in radian, into two clusters using a k-means (k=2) clustering. After empirical investigation, we noticed that an epidermal cell touches two cortex cell file when the distance between the two clusters is bigger than 0.4 radian and therefore is a trichoblast cell, otherwise the cell is an atrichoblast cell.

## Acknowledgements

We thank Prof. Peter Schopfer, Dr. Patrick Laufs, Nicolas Arnaud and the members of our teams for their critical discussion and reading of the manuscript. We also gratefully acknowledge the excellent technical support from Roland Nitschke (Life Imaging Centre, ZBSA, Freiburg). This work was supported by the Collaborative Research Center 746, the Excellence Initiative of the German Federal and State governments (EXC 294), Bundesministerium für Bildung und Forschung (BMBF) (FKZ 0315329B, 0101-31P5914) and Deutsches Zentrum für Luft und Raumfahrt. We finally want to gratefully acknowledge EMBO for the long-term postdoctoral fellowship awarded to T.B.

